# Phytoplankton phenology through gene expression during the North Atlantic spring bloom decline

**DOI:** 10.1101/2025.10.15.682395

**Authors:** Meredith G. Meyer, Olivia Torano, Natalia L. Llopis-Monferrer, Nicolas Cassar, Melanie R. Cohn, Mark A. Brzezinski, Adrian Marchetti

**Affiliations:** Department of Earth, Marine, and Environmental Sciences, University of North Carolina at Chapel Hill, Chapel Hill, NC, USA; Monterey Bay Aquarium, Research Institute, Moss Landing, CA, USA; Sorbonne University, CNRS, UMR7144 Adaptation and Diversity in Marine Environment (AD2M) Laboratory, Ecology of Marine Plankton team, Station Biologique de Roscoff, Roscoff, France; Division of Earth and Climate Sciences, Nicholas School of the Environment, Duke University, Durham, NC, USA; Department of Ecology, Evolution and Marine Biology, Marine Science Institute, University of California, Santa Barbara, CA, USA

**Keywords:** Phytoplankton, Spring bloom, Metatranscriptomics, North Atlantic

## Abstract

While phytoplankton dynamics in the annual North Atlantic spring bloom have been well characterized, the physiological underpinnings driving these changes and their net impact on the biogeochemistry of the region are less understood. Phytoplankton metabolism is both affected by, and influences the region’s nutrient cycling, primary production, and ultimately, the fate of carbon export. Thus, developing an understanding of these processes is critical. Phytoplankton biomass, biological rates, and gene expression data along with associated environmental parameters were measured as part of the NASA EXport Processes in the Ocean from RemoTe Sensing program’s campaign to the North Atlantic to evaluate the relationships amongst these processes within the four most dominant phytoplankton groups (diatoms, dinoflagellates, haptophytes, and chlorophytes) during the spring bloom. We observe a transition from a period dominated by active diatom growth (defined as Phase I) to a period dominated by non-diatom phytoplankton groups (Phase II). Silicic acid depletion appears to limit overall production and reduce competition from diatoms, likely leading to enhanced contributions of haptophytes in Phase II. Expression of key protein-encoding genes involved in cell maintenance, photosynthesis, and nitrogen and vitamin metabolisms varied amongst the taxa throughout the observation period. Expression patterns of diatom genes involved in silicon transport suggest an apparent uncoupling between genes involved in nitrate uptake and photosynthesis, resulting in an increase in silicification independent growth. Our analysis demonstrates the utility in combining gene expression with biological rate processes to provide a more holistic view of phytoplankton bloom dynamics and phenology.

## 1. Introduction

Due to their substantial contribution to seasonal carbon export, spring blooms in the North Atlantic have been extensively studied (Henson et al., 2019; Buesseler et al., 2020; Johnson et al., 2024), with particular focus on drivers such as physical processes, nutrient concentrations, phytoplankton community composition, and seeding of benthic communities (Hartman et al., 2010; Behrenfeld 2010; Lampitt et al., 2010; Henson et al., 2012). However, substantially less focus has been paid to the specific physiological mechanisms that drive phytoplankton bloom succession and phenology and how these mechanisms are simultaneously linked to the physical and biological drivers of community composition, and the net impacts on biogeochemical cycling during the bloom period. Recent studies have shown that the oceans are experiencing a new form of succession, a taxonomic shift in phytoplankton community composition toward smaller-celled groups, making the oceans overall “greener” (i.e., a shift toward predominance of green algal species; Henson et al., 2021; Cael et al., 2023). Therefore, a better understanding of how differences in physiology between phytoplankton groups in response to environmental dynamics impacts net changes in energy flow and carbon cycling is critical for predicting ecosystem functioning and biogeochemical processes.

The annual North Atlantic Spring bloom appears to be initiated in late winter to early spring when a combination of increased irradiance and fluctuating mixed layer depths entrain nutrients from depth and recouple plankton populations in space, causing increased phytoplankton growth and production beginning around March/April (Sverdrup, 1953; Behrenfeld 2010; Hartman et al., 2010). The dominant phytoplankton group during the initiation of the bloom are frequently diatoms which hold a competitive advantage due to their enhanced nutrient acquisition and storage abilities and fast growth rates under high nutrient conditions (Leynart et al., 2004; Marchetti et al., 2006). As the bloom begins to decline, the phytoplankton community shifts towards dinoflagellates and haptophytes (Henson et al., 2012; Sundby et al., 2016). However, recent studies have shown that the proportion of diatoms does not decline as much as previously thought, despite changes in their rates of primary production (Meyer et al., 2024).

Physiological differences between phytoplankton groups make relating taxonomy to physical and biogeochemical parameters important for our understanding of large-scale ecosystem dynamics. Key differences exist in the traits among diatoms, dinoflagellates, and haptophytes including nutrient requirements, stoichiometry and metabolism (Garcia et al., 2018). In addition to all autotrophs’ requirement for carbon, nitrogen, phosphorus, and micronutrients, diatoms have a cellular requirement for silicic acid, and many abundant haptophytes (coccolithophores) require calcium carbonate, creating additional constraints on growth and primary production. Diatoms and haptophytes have been shown to primarily engage in new production (i.e., primary production that utilizes “new” nitrogen sources to the euphotic zone such as nitrate and dinitrogen gas; Dugdale and Goering, 1967). This is believed to support shorter food webs (Michaels and Silver, 1988) and promote enhanced carbon export efficiency (Eppley and Peterson, 1979). However, all phytoplankton groups have varying cell sizes, carbon content, sinking rates, photosynthetic efficiencies, and mortality rates, undoubtedly impacting their contribution to carbon export and sinking flux.

Here, we examine changes in environmental conditions that drive taxonomic and inferred physiological shifts during the decline phase of the North Atlantic spring bloom at the molecular level and the manner in which these changes relate to net bloom and biogeochemical changes during the NASA EXport Processes in the Ocean from RemoTe Sensing (EXPORTS) campaign to the Porcupine Abyssal Plain (PAP) region in the North Atlantic. The EXPORTS campaign was initiated to compare and contrast the drivers and mechanisms of carbon export in a low productivity, low export region (Ocean Station Papa; Siegel et al., 2021) in the subarctic North Pacific to a high productivity, high export region in the North Atlantic (PAP; Johnson et al., 2024). This study focuses on the campaign to the PAP region in May of 2021 where an anticyclonic eddy was occupied for 31 days. The comprehensive nature of the program enabled the integration of phytoplankton transcriptomic data with nutrient concentrations, rates of primary production, phytoplankton community composition within the mixed layer, and estimates of carbon export. This allowed us to evaluate how phytoplankton metabolic processes within distinct groups, as inferred through their gene expression, changed during the bloom decline.

## 2. Materials and Methods

### 2.1 Sampling strategy

Samples for primary production, phytoplankton biomass, environmental parameters, and RNA were collected between Yeardays (YDs) 126-148 (May 6^th^ – 28^th^) in 2021 within the vicinity of the PAP site (49.0°N, 16.5°W) in the North Atlantic Ocean (Fig. 1). Sample collection occurred aboard the Royal Research Ship (*RRS*) James Cook. The ship sampled in a quasi lagrangian pattern according to temperature and salinity classifications from a profiling float drogued at 100 m. Mixed layer depths (MLDs) were determined according to the 0.03 kg m^-3^ density differential of CTD casts aboard the RRS Cook. Samples for dissolved inorganic nutrients, including nitrate (NO_3_ ^-^), phosphate (PO_4_ ^3-^), and silicic acid (Si(OH)) and biogenic silica (BSi) were collected throughout the euphotic zone as according to Meyer et al. (2024) and Brzezinski et al. (2024). For a more detailed discussion of MLD delineation and sampling schemes, see Johnson et al. (2024) and Meyer et al. (2024). All production, biomass, and environmental data have been deposited in the SeaWiFS Bio-optical Archive and Storage System (SeaBASS; https://seabass.gsfc.nasa.gov/cruise/EXPORTSNA/). Raw sequences are deposited at the National Center for Biotechnology Information (NCBI; ascension number PRJNA1072555).

**Figure 1.**
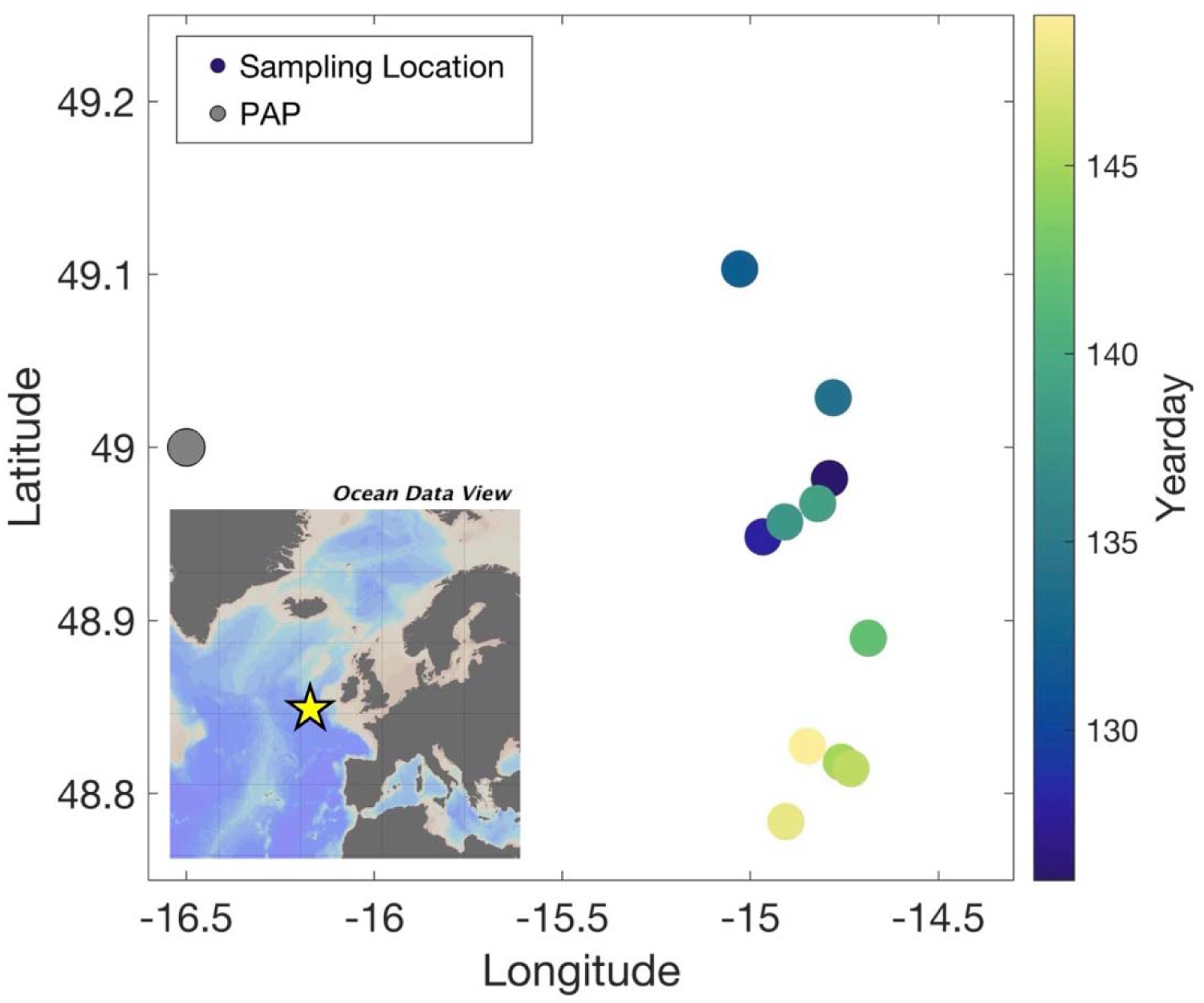
EXPORTS productivity sampling sites in the North Atlantic Ocean. Map of sample locations during the North Atlantic EXPORTS field campaign collected aboard RRS James Cook. Symbol color corresponds to yearday of sampling. The Porcupine Abyssal Plain site is noted in gray. The location of the PAP site and our sample sites in relation to the European coast are indicated by the star in the inset map.

### 2.2 Phytoplankton biomass

Triplicate 400 mL bottles were collected for chlorophyll *a* (Chl *a*) concentrations from the depths corresponding to those for the primary production experiments (see below). Bottles were size-fractionated as according to Meyer et al. (2024), resulting in a < 5 µm size-fraction (small cells) and a ≥ 5 µm size-fraction (large cells) that were summed to produce a total, whole water estimate. Once collected, filters were immediately frozen at -20°C. Samples were run on a Turner 10AU fluorometer aboard the *RRS* Cook via the method described in Graff and Rynearson (2011). Estimates of total and size-fractionated (<5 µm, ≥5 µm) particulate carbon (PC) and particulate nitrogen (PN) were provided from the primary production samples (Meyer et al., 2024).

### 2.3 Primary production

On days when the MLD was deeper than the euphotic zone (defined as the 1% irradiance (I_o_) level), 1 L bottles were collected from five depths corresponding to 65%, 20%, 10%, 5%, and 1% I_o_. On days when the MLD was shallower than the euphotic zone, the 1 L bottles sampled were confined to the I_o_ levels within the mixed layer, which was typically 2-4 depths.

Triplicate bottles were inoculated with 180 µmol L^-1^ of NaH^13^ CO_3_^-^ isotope, approximating an 8% addition relative to ambient dissolved inorganic carbon (DIC) concentrations and 0.2-0.7 µmol L^-1^ of Na^15^NO_3_ ^-^ isotope, approximating 16% ambient NO_3_ ^-^ concentrations (targeted as <10% of ambient concentrations). Bottles were incubated in on-deck, flow-through incubators at their respective light levels for 24 hours before filtration. Estimates of net primary production (NPP) were made from DIC uptake rates (Marra, 2002; Marra, 2009; Barber and Hilting, 2007) whereas ^15^NO_3_ ^-^ uptake rates were converted to carbon units via the Redfield ratio and used to approximate new production (Dugdale and Wilkerson, 1986; Meyer et al., 2024). Triplicate bottles provide size-fractionated (<5 µm, ≥5µm, and total) samples. Estimates of siliceous phytoplankton specific contributions to total NPP were estimated via silicon-32 uptake (ρSi) according to Brzezinski et al. (2024). Integrated mixed layer values were calculated according to trapezoidal integration (Meyer et al., 2024).

### 2.5 RNA collection and extraction

Seawater was collected in triplicate from surface (∼5 m) Niskin bottles via a standard CTD rosette. After collection, seawater was immediately pumped via low peristaltic pressure for a maximum of 40 minutes through a 0.8 µm Supor filter. This resulted in an average of 6 L being filtered. RNA filters were placed into 5 mL cryovials, flash frozen in liquid nitrogen and stored at -80°C until onshore extraction.

RNA was extracted using the Qiagen RNAqueous-4PCR plant tissue extraction kit as according to manufacturer instructions with modifications to enable extraction from large filters (142 mm). These were as follows: filters were cut into small pieces into a 50 mL centrifuge tube. Glass beads (0.5 mL) and 7 mL of lysis buffer were then added and vortexed on high speed. Filter pieces were removed and the lysis buffer solution was passed through a filter column and eluted. Extracted RNA was treated with DNAse, underwent cleanup via a RNeasy MinElute Cleanup kit, and were stored at -80°C until sequencing.

### 2.6 RNA sequencing and metatranscriptomic analysis

Extracted RNA was sequenced at Azenta Life Sciences via Illumina HiSeq on two lanes with ∼350M raw paired-end reads per lane and single index, 2 x 150 bp per lane.

Paired-end reads underwent trimming via TrimGalore (v.0.6.2; Altos Labs) to remove sequence adapters and poly-A tails followed by FastQC (v.0.11.9) for quality control, removing any reads that were deemed of too low quality for the analysis (quality score < 20 or < 50 bp). Trinity (v.2.8.6) was used for de novo assembly of reads according to individual sample day.

Following Trinity assembly, cdhit (v.4.8.1) was used to combine individual assemblies into a singular grand assembly (Fu et al., 2012). The grand assembly was annotated, blasting via Diamond, for best hit functional annotation (e-value cut off = 10^-5^) via the Kyoto Encyclopedia of Genes and Genomes (KEGG; Release 88.2) and taxonomic annotation via PhyloDB (github.com/allenlab/PhyloDB). Additional, manual blasting of the grand assembly for certain sentinel genes of interest occurred. Fasta files for 41 silicon metabolism-related genes of interest are described in Durkin et al. (2012), Marron et al. (2016), and Durak et al. (2016; Table S1). End-to-end read alignment occurred via Salmon (v.10.9.1; Patro et al., 2017) with aligned files and annotated grand assemblies being combined via tximport (Soneson et al., 2015) and resulting in read counts per million (CPM).

Two versions of the dataset were visualized. The first version includes gene CPMs normalized to the datasets first separated into four dominant phytoplankton taxonomic groups (diatoms, dinoflagellates, haptophytes, and chlorophytes). This normalization provided an examination of the gene expression changes within a taxonomic group throughout the observation period. The second version had CPMs normalized to the whole community dataset, providing a community-wide comparison of changes in gene expression among the taxonomic groups throughout the observation period. Additionally, physiologically useful ratios of gene CPMs and rates were evaluated.

## 3. Results

### 3.1 Environmental and phytoplankton parameters

Sampling days were separated into Phase I (PI; YDs 126 and 128) and Phase II (PII; YDs 132-149) based on the impact from a storm event which occurred from YDs 127-130 and substantially altered the physical, chemical and biological properties of the site, including mixed layer depth, nutrient inventory, and phytoplankton dynamics (Johnson et al., 2024; Meyer et al., 2024). Two additional storms occurred, but the first storm had the largest impact on mixed layer properties. This separation is supported by the averaged differences by phase, where phase- averages in MLD, nutrient concentrations (NO_3_ ^-^, PO_4_ ^3-^, Si(OH)), NPP, new production, proxies for the contribution of the large size-fraction to phytoplankton biomass (Chl *a*, PC, PN), BSi, and ρSi show substantial differences (Fig. 2). Partially due to the higher number of samples (8 in PII vs. 2 in PI), variability within PII datasets is much larger, as exemplified by the larger variability and variety of patterns between temporal data points (Fig. 2). Overall, macronutrient concentrations are higher (28.9%, 38.0%, 88.4% for NO_3_^-^, PO_4_^3-^, and Si(OH)_4_, respectively) in PII. As suggested by the percent increase, the most significant change (p < 0.05, Wilcoxon rank sum test) is in mixed-layer Si(OH)_4_ which averaged 0.1 ± 0.11 µg L^-1^ in PI and 1.12 ± 0.63 µg L^-1^ in PII in the mixed layer. ρSi exhibited a similar increase in PII to the macronutrients.

**Figure 2.**
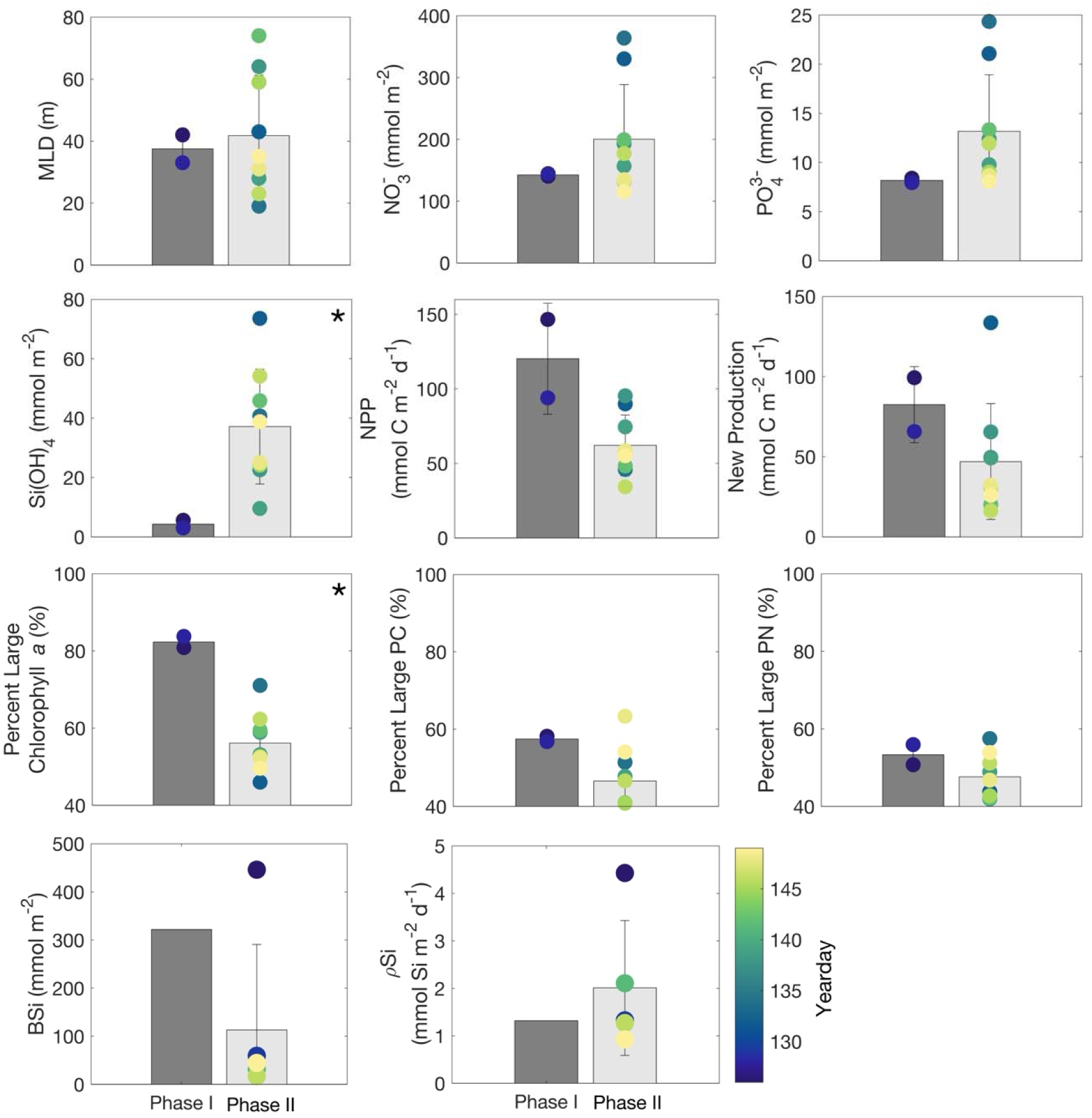
Bar plots of average mixed layer depth (MLD; m) and mixed layer integrated nitrate concentration (NO_3_^-^; mmol m^-2^), phosphate concentration (PO_4_^3-^, mmol m^-2^), silicic acid concentration (Si(OH)_4_; mmol m^-2^), net primary production (NPP; mmol C m^-2^ d^-1^), new production (mmol C m^-2^ d^-1^), and relative contribution of large phytoplankton (>5 μm) to chlorophyll *a* (Chl *a;* %), particulate carbon (PC; %) and particulate nitrogen (PN; %), biogenic silica (BSi; mmol m^-2^) and silicic acid uptake (ρSi; mmol Si m^-2^ d^-1^) in Phase I (dark grey bars) vs. Phase II (light grey bars). Error bars correspond to standard deviations per sample phase. Symbol colors correspond to yearday. Asterisks indicate a statistically significant relationship based on Wilcoxon rank sum testing.

Conversely, NPP, new production, the contribution of large phytoplankton to Chl *a*, PC, and PN, and BSi are all higher in PI. There is a large and statistically significant difference (p < 0.05, Wilcoxon rank sum test) in the relative contributions to Chl *a* by size-fractions between PI and PII, where the large size-fraction accounts for 82.3 ± 2.0% in PI but drops to 56.1 ± 7.7% in PII. The substantial differences between phases resulted from a net deepening of the MLD (average of 32.8 m in PI and 42.6 m in PII) which caused horizontal and vertical advection of biomass and an injection of nutrients from below (Johnson et al., 2024).

### 3.2 RNA sequencing statistics

Metatranscriptome sequencing depth ranged from 18.5 – 33.1 million reads with an average of 27.8 million reads per sample (Table S2). GC content was relatively consistent ranging from 62.0 – 68.4% with an average of 65.9 ± 1.64% (Table S2). Statistical analysis of sequenced reads show noteworthy differences between sampling days grouped into PI versus those in PII (Fig. 3), supporting our separation of the samples into these two groups.

**Figure 3.**
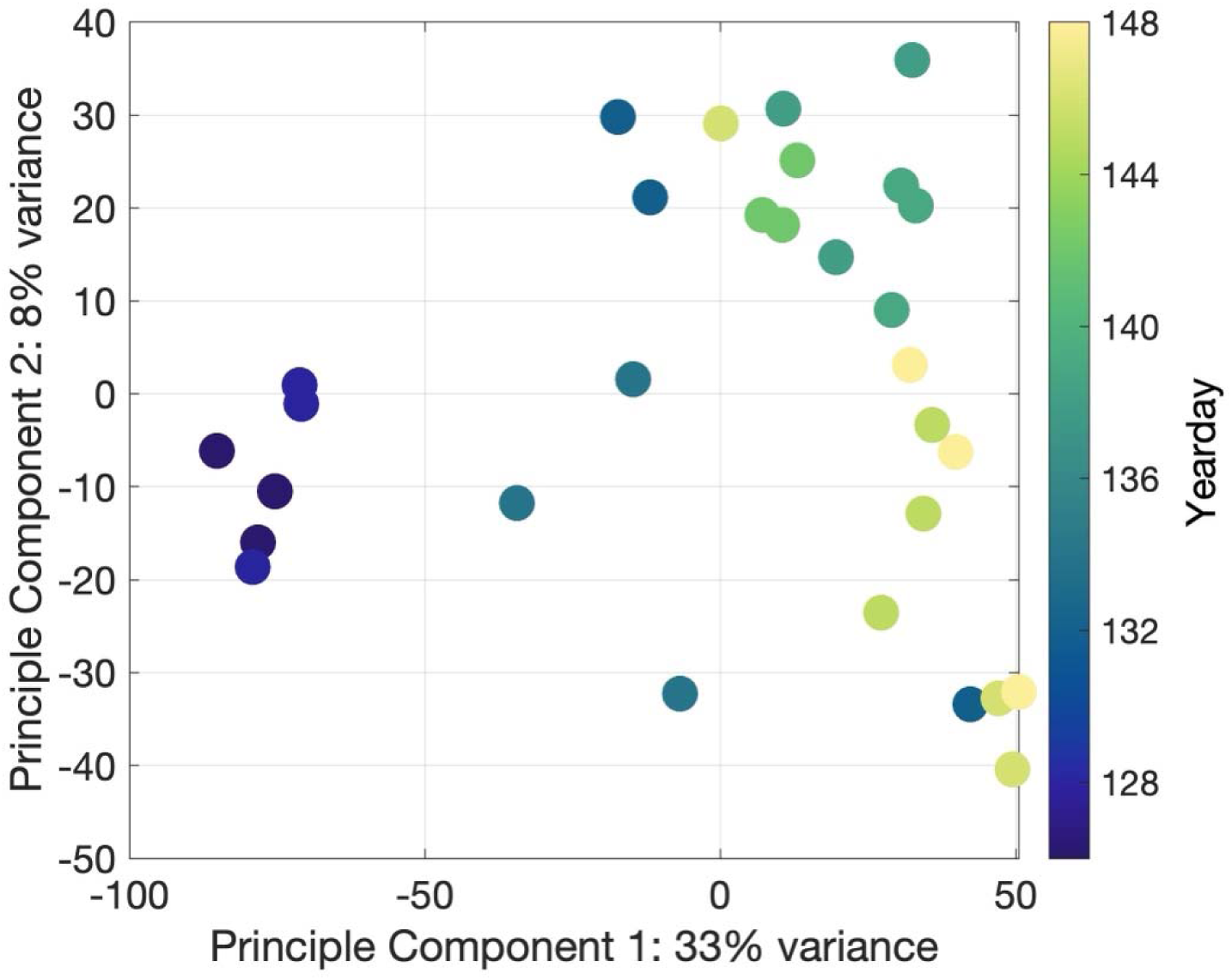
Principle component analysis of metatranscriptomic sequence samples (n=30). Samples are colored by yearday.

The total number of reads ranged from 1.85 x 10^7^ on YD 126 to 3.31 x 10^7^ on YD 134 (Fig 4A). Diatoms constituted the largest percent of reads (approximately 58%) during Phase I, but decreased in Phase II where haptophytes began to constitute the largest relative percentage (46-58%; Fig 4A). Both phytoplankton groups exhibited substantial variability with diatoms decreasing 47%, and haptophytes increasing 47% over the observation period. The decline in diatom reads from Phase I to Phase II and the proportional increase in haptophyte contribution to total contigs is noteworthy, supporting both a change in net biomass contribution that coincides with a decline in production as well as a shift in community composition consistent with what was observed in other EXPORTS datasets (Meyer et al., 2024). Dinoflagellates and chlorophytes remained relatively consistent at approximately 8% and 18%, of the read proportions, respectively.

**Figure 4.**
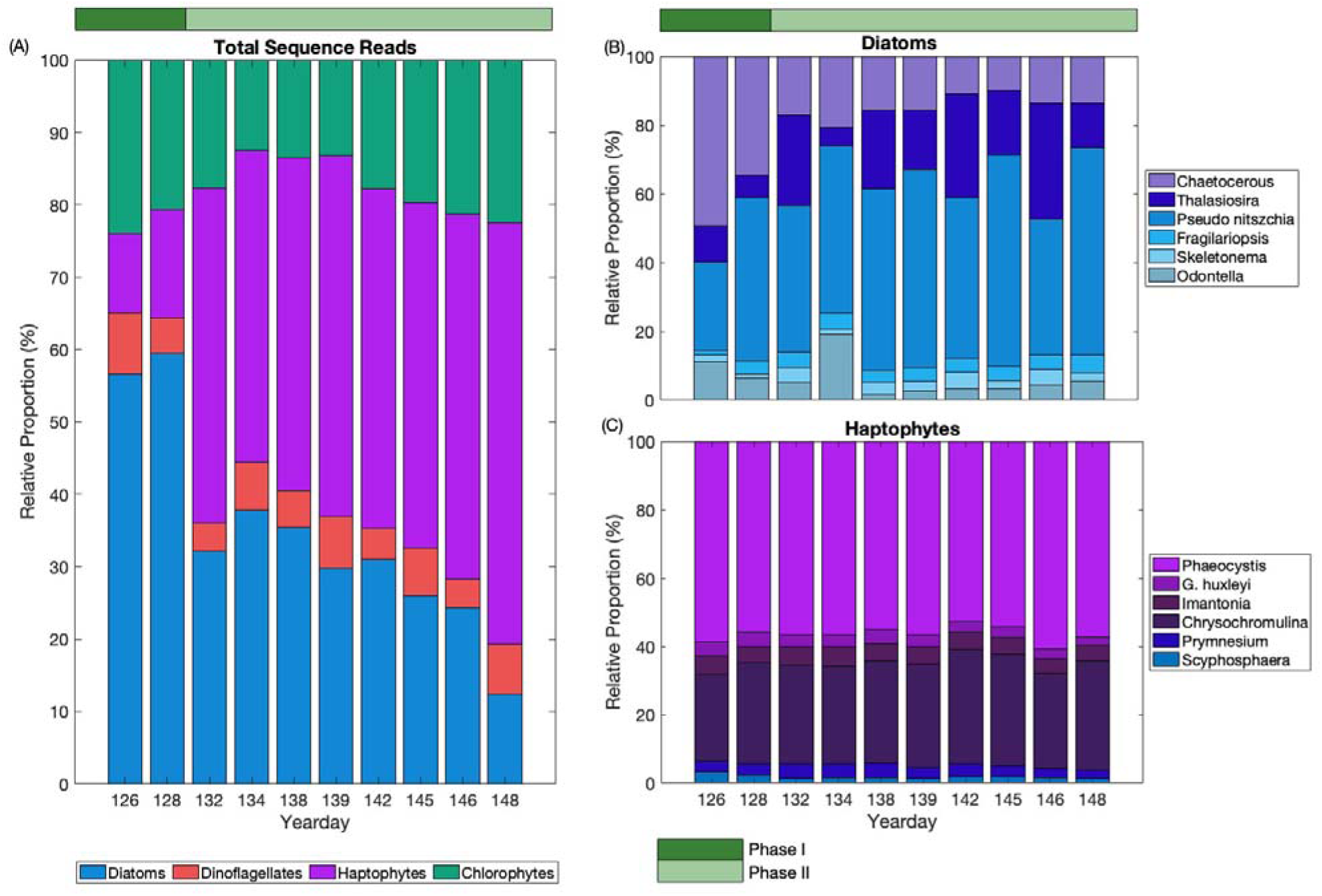
(A) Total relative proportion (%) of the four most abundant phytoplankton groups based on RNA sequence reads, and (B,C) the relative proportion (%) of the most abundant genera in diatoms and haptophytes based on sequence reads. *G. huxleyi* is *Gephyrocapsa huxleyi* (formerly *Emiliania huxleyi*). Horizontal green bars indicate phase.

### 3.3 Taxonomic representation

Within the diatoms, members of *Pseudo-nitszchia* (48%), *Chaetocerous* (20%), and *Thalassiosira* (18%) were the three most represented genera based on RNA sequence reads (Fig. 4B). The haptophytes had a similar predominance of one main genera with *Phaeocystis* accounting for 56% of total reads and *Chrysochromulina* and *Prymnesium* accounting for 30% and 4%, respectively (Fig. 4C; see Supplemental Results S2.1 for dinoflagellate and chlorophyte representation).

### 3.4 Transcript abundance trends

The transcript abundance of key genes can be categorically separated into those relating to nitrogen metabolism, photosynthesis, vitamin metabolism, and for diatoms, silicon metabolism. Reads are presented in two ways: normalized both within phytoplankton groups (Fig. 5A) and to the whole community (Fig. 5B). Normalizing by whole community transcripts provides a comparative analysis on how the proportions of each metabolic activity are changing in each phytoplankton group relative to the other groups whereas the taxon-specific normalization provides an estimate of changes in gene expression within a phytoplankton group. Despite substantial variability amongst phytoplankton groups, distinct patterns are visible between how the expression of various genes changed through time, particularly between PI and PII.

**Figure 5.**
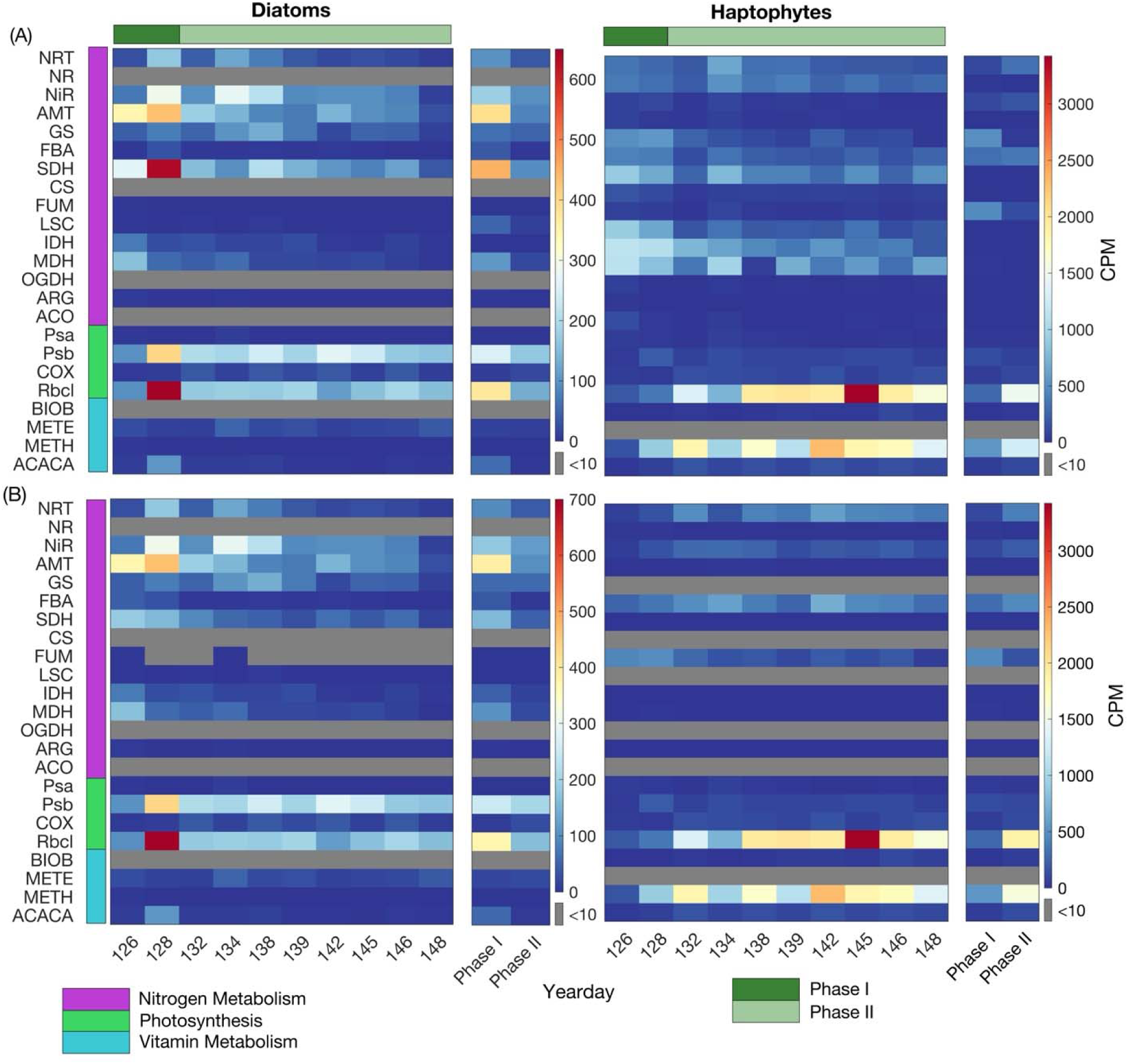
Heatmaps of transcript counts per million (CPMs) of key nitrogen metabolism, cell maintenance, hotosynthesis, and vitamin metabolism genes (indicated by left hand bars) for diatoms and haptophytes with counts normalized within each taxonomic group (A) and to the whole community (B). Plots are by yearday nd phase average (normalized the number of sample days) with the color bar indicating CPM for each gene. orizontal green bars indicate phase. Full gene names are: nitrate transporter (*NRT*), nitrate reductase (*NR*), nitrite transporter (*NiR*), ammonium transporter (*AMT*), glutamine synthase (*GS*), flavodoxin (*FBA*), succinate dehydrogenase (*SDH*), citrate synthase (*CS*), fumarate (*FUM)*, succinyl-CoA synthetase (*LSC*), socitrate dehydrogenase (*IDH*), malate dehydrogenase (*MDH*), oxoglutarate dehydrogenase (*OGDH*), arginase (*ARG*), aconitase (*ACO*), photosystem I (*Psa*), photosystem II (*Psb*), cytochrome c oxidase (*COX*), RUBisCO (*Rbcl*), biotin synthase (*BIOB*), methionine synthase (*METE*), methyltransferase (*METH*), and cetyl-CoA carboxylase (*ACACA*).

Diatoms displayed the highest expression of all genes on YD 128 where genes related to photosynthesis exhibited their maximum transcript abundances of the entire observation period (Fig. 5A). Within the diatom-specific normalization, genes encoding for photosynthetic genes, RubisCO (*Rbcl*) and photosystem II (*Psb*), and nitrogen assimilation genes, ammonium transporter (*AMT*), nitrite reductase (*NiR*), and acetyl-CoA carboxylase (*ACACA*) increased 5.8-, 3.1-, 0.2-, 3.1-, and 9.4-fold change in PI from YD 126 to YD 128, respectively. The changes in counts normalized to the entire community are comparable (Fig. 5B). Of the seven diatom-specific silicon transport genes analyzed, two (Si4 and Si7) showed similar trends to the photosynthesis related genes, peaking on YD 128 (Fig. 6B; 6C). These genes are annotated as silicon transporters from *Fragilariopsis cylindrus* and *Phaeodactylum tricornutum*, respectively (Table S1; Mock et al., unpublished; Bowler et al., 2008). YD 128 coincides with the highest number of sequences of the four primary taxa for any sample day as well as the highest proportional contribution of diatom sequences to total sequences (35.9%; Fig. 4A). During PII, transcripts of *Psb*, *Rbcl*, and *NiR* were on average higher than on YD 126, a noteworthy distinction, suggesting that despite lowered rates, NPP and new production still occurred (Fig. 2; Fig. 5A, 5B). The second highest counts for Si4 and Si7 are also observed in PII on YD 134 (Fig. 6B; 6C), which coincides with the beginning of the second large storm event and an increase in Si(OH)_4_ concentrations in the mixed layer (Johnson et al., 2024).

**Figure 6.**
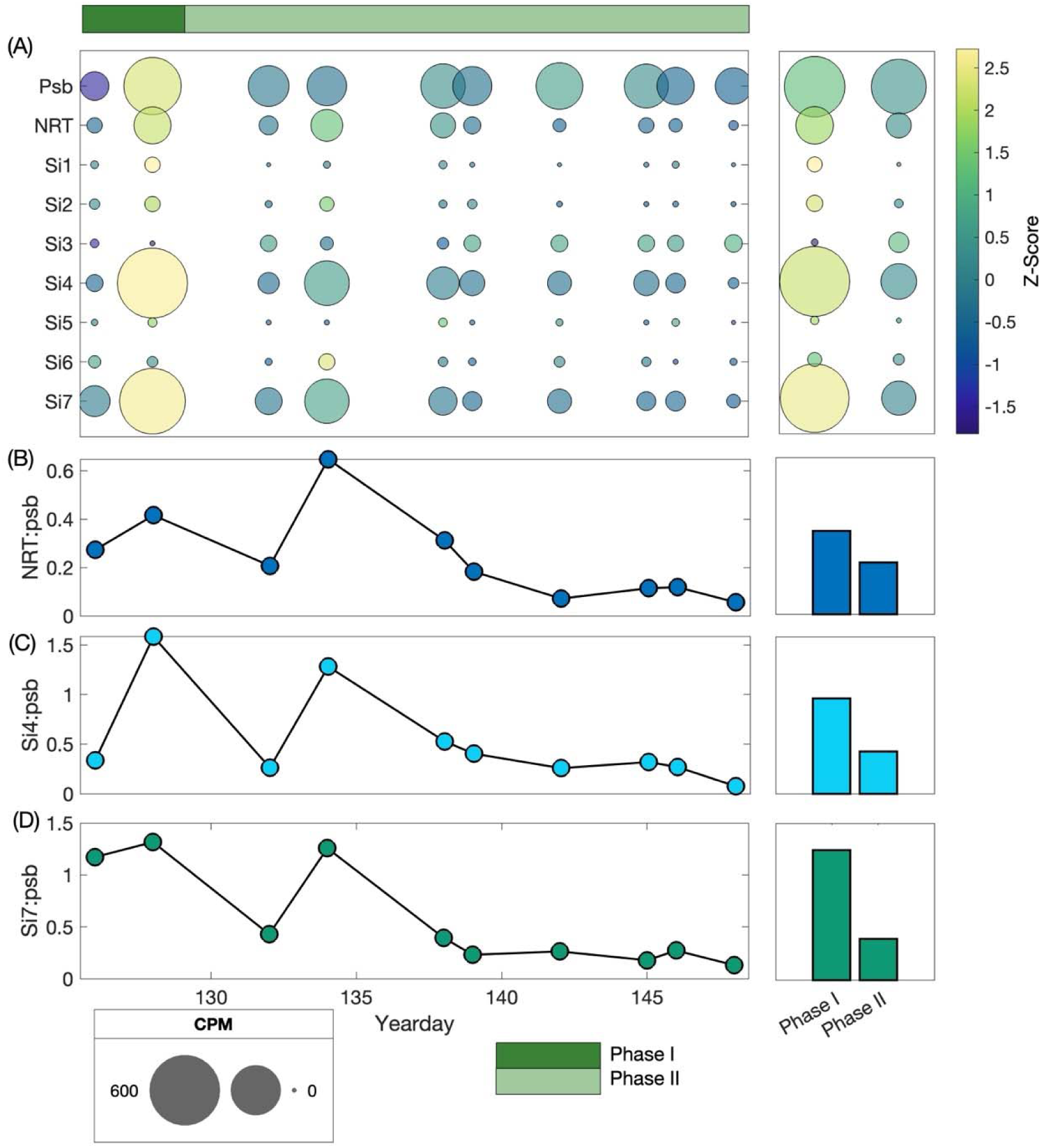
(A) Transcript counts per million (CPMs) of diatom *Psb*, *NRT* and silicon transport genes normalized within the diatom group by yearday and phase average normalized to number of sample days and ratios of diatom (B) *NRT:Psb*, (C) Si4:*Psb*, and (D) Si7:*Psb* by yearday and phase average. The size of the symbol in panel (A) corresponds to CPM. The color bar indicates the Z-score. Further information on the Si gene4 and Si gene7, both of which are silicic acid transporters, is available in Table S1. Green bars indicate phase.

Haptophytes appear to exhibit the most appreciable difference between gene expression patterns normalized at the phytoplankton group versus whole community level of any of the examined taxa (Fig. 5A,B). This is apparent from numerous genes involved in cell maintenance and nitrogen metabolism which are better represented in the haptophyte specific normalization but are weakly expressed (CPM <10) in the community normalization, including glutamine synthase (GS), citrate synthase (CS), isocitrate dehydrogenase (IDH), oxoglutarate dehydrogenase (OGDH), and aconitase (ACO) (Fig. 5B). However, some consistent patterns still emerge between the two approaches and suggest substantial differences exist between PI and PII, particularly in terms of photosynthesis. In the haptophyte specific normalization, *Rbcl* and *METH* are on average 5.8- and 2.0-fold higher in PII with similar patterns for the community normalization. While the differences between phases are not as obvious, *NRT* also increased substantially (>1400%) between PI and PII. Across both PI and PII, *Psb* exhibited higher abundances than *Psa*, suggesting that like in diatoms, haptophytes may be engaging in NO_3_- dependent primary production (i.e., new production) with a preference for expression of *Psb* (Fig. 5A, 5B). Similar to diatoms, these changes between phases coincide with shifts in the relative proportion of haptophyte to total sequences (Fig. 4B). Contrastingly, gene expression patterns in dinoflagellates and chlorophytes were consistent across phases (see Supplemental Results 2.2).

### 3.5 Gene expression and nutrient uptake patterns

To gain a better understanding of our gene expression data relative to our biogeochemical rate data, we calculated numerous gene ratios involving key genes for overall primary production and diatom cell growth, including *NRT*:*Psb* and Si:*Psb* (Fig. 6; S5). For diatoms, the Si-related gene ratios were assessed and compared to silicic acid uptake (^32^Si-*ρ*Si) rates obtained during EXPORTS. Ratios of Si4:*Psb*, Si7:*Psb*, Si4:*NRT*, and Si7:*NRT* were all higher in PI relative to PII (Fig. 6B; 6C). Overall, ratios with Si7:*Psb* and Si7:*NRT* during PI were higher than ratios of Si4:*Psb* and Si4:*NRT* despite the average Si4 transcript abundance being higher (137 CPMs) and exhibited more temporal variability (standard deviation of 187) than Si7 (average = 129 ± 154). When evaluating *ρ*Si:*ρ*DIC vs. Si4(7):*Psb* and *ρ*Si:*ρ*NO_3_ ^-^ vs. Si4(7):*NRT* using a principle component analysis, the strongest relationships were between large size-fraction *ρ*Si:*ρ*DIC vs. Si7:*Psb* (R^2^ = 0.65) and large size-fraction *ρ*Si:*ρ*NO_3_ ^-^ vs. Si7:*NRT* (R^2^ = 0.51). Both relationships were negative (Fig. S5).

## 4. Discussion

### 4.1 Spring bloom phenology

The annual North Atlantic spring bloom is known to exhibit a predictable taxonomic succession from diatom to haptophyte dominance coincident with a decline in nutrient concentrations, particularly NO_3_^-^, and resulting phytoplankton biomass (Henson et al., 2012). Here, our findings suggest that the 2021 spring bloom exhibited a typical taxonomic succession associated with bloom decline. However, instead of NO_3_^-^ limitation, Si(OH)_4_ depletion in Phase I may have created an unexpected physiological response in diatoms that hindered primary production yet resulted in enhanced pulses of carbon export (Daniels et al., 2015). Our transcriptomic results support this finding, suggesting this physiological response persists into Phase II when Si(OH)_4_ concentrations were partly replenished with storm-induced mixing.

Silicic acid limitation of diatoms was confirmed by Si addition incubation experiments (Brzezinski et al., 2024), but the inability for diatoms to recover physiologically in the days to weeks following natural Si(OH)_4_ replenishment is noteworthy and contrary to previous findings (De La Roche and Passow, 2004). Combined with gene expression trends, this suggests a decoupling between intracellular activity and ecosystem rate measurements due to substantial reductions in the biomass of large cells such as diatoms, likely caused by loss processes such as grazing and export.

As is evident through the transcript abundance plots (Fig. 5; Fig. 6), our analysis suggests strong temporal differences between the physiological patterns amongst the different phytoplankton groups, particularly diatoms and haptophytes. For diatoms, YD 128 clearly represents a transition where the decoupling between the transcription of key metabolic genes (*Rbcl*, *Psb*, *AMT*, and *NRT*), NPP and new production rates is most pronounced. YD 128 marks the peak in photosynthesis gene expression, but this does not appear to translate into higher NPP or new production. For haptophytes, YD 126 was the most distinct (Fig. S3; See Supplemental Methods S1.1. and Results S2.3.) and represents a day of comparatively lowered expression of genes related to new production (*NRT*:*Psb* is 1.64; Fig. S5). For haptophytes, YD 126 marks the lowest transcription of *Psb* and *METH* with 20 and 755 fewer transcripts relative to peak values of 210 and 2302 on YD 128 and 142, respectively. It is noteworthy that for both the diatoms and haptophytes, their day with the largest difference in transcripts abundances occurs in PI. This reinforces the observation that the transition from PI to PII marks a substantial shift in the biogeochemical and taxonomic attributes of the system coincident with the first mixing event.

### 4.2 Phytoplankton succession during bloom decline

To better understand the drivers of these taxonomic and temporal differences, key phase dynamics were examined, paying particular attention to changes in the Si-related genes relative to changes in better understood metabolic genes, such as *Psb*, *NRT*, etc. The two most expressed Si genes, Si4 and Si7, have strong similarity to known silicic acid transporters (*SIT*) in the NCBI repository (Table S1; Mock et al., unpublished; Bowler et al., 2008). However, there is still a general lack of understanding of silicic acid transporter activity and silica dynamics in diatoms, partially due to the poorly conserved nature of *SITs*, which allows them to quickly adapt to changes in their environment (Brembu et al., 2017). Gene expression analysis has shown that some *SITs* cluster together, suggesting coordinated activity (Thamatrakoln and Hildebrand, 2005). The similarity of patterns in Si4 and Si7 may be an example of this type of clustering. Whether these SITs can act independently or require co-expression is uncertain.

Previous studies have shown how *SIT* activity can be inversely related to Si(OH)_4_ concentration (Hildebrand et al., 1998; Oh et al., 2018) but are closely related to the cell cycle (Brembu et al., 2017). While some uncertainty exists around the subject (Maniscalco et al., 2023), enhanced expression of *SITs* without a congruent increase in primary production may support the idea that this could lead to enhanced silicification, creating heavier ballasted diatom cells that may facilitate faster sinking rates and enhanced carbon export out of the euphotic zone (Finkel, 2016). This is consistent with optical and sediment trap data which indicated more large particles were exported from the upper water column and a higher representation of biogenic silica in trap material in the latter part of the observation period (Siegel et al., 2024; Brzezinski et al., 2024).

While overall nutrient uptake rates in PI were driven by diatoms, these dynamics appear driven mostly by non-diatoms in PII, particularly haptophytes. Despite having fewer overall transcripts, due to smaller genomes in some species (Sanchez Puerta et al., 2005), haptophytes may be a key contributor to primary production in PII of the observation period when rates of NPP and new production have declined (Fig. 2; Table S3). During this time, haptophytes maintain substantial transcription of key genes for photosynthesis (*Rbcl*) and methionine synthase (*METH*) as well as a high (average = 4.77 ± 1.89) ratio of *NRT*:*Psb*, potentially suggesting utilization of NO_3_ ^-^ (Fig. 5A, 5B; Fig. 8A; Sunda and Huntsman, 1995), consistent with the idea of reduced resource competition during this period. This idea contradicts the previously proposed rising tide hypothesis (i.e., conditions favorable for diatoms will be likewise favorable for other phytoplankton groups, and vice versa; Brown et al., 2008) and supports the notion that resource competition may be lowered during this time as diatoms are less metabolically active. The high proportion of *Phaeocystis*, which contain a nitrogen rich colonial fluid (Arrigo et al., 1999; Hales and Takahashi, 2004) is noteworthy and raises the question of potential luxury storage of nutrients and competitive advantage.

In our two less dominant phytoplankton groups, dinoflagellates and chlorophytes, we do not see as strong gene expression trends as in diatoms and haptophytes. One exception to this is *Psb* in dinoflagellates which substantially increases in abundance from PI to PII (Fig. S2A; S2B). This increase coincides with a substantial decrease in mixed layer biomass (46.6% in Chl *a* and 23.4% in PC) and may signify a shift toward greater autotrophy within the dinoflagellate community. Additionally, the dinoflagellate community composition does not appear to change substantially (Fig. S1A), perhaps suggesting the shift may result from mixotrophic capabilities rather than a shift in the taxonomy (Jones et al., 2025). This notion is further supported by the finding that the three most transcriptionally active, representative dinoflagellate genera, *Karlodinium, Alexandrium, and Karenia*, are all known to be mixotrophic (Adolf et al., 2006; Lee et al., 2016; Glibert et al., 2009). As a much smaller proportion of the phytoplankton community, chlorophytes key metabolic genes, including *Rbcl*, *Psb*, NRT, AMT, etc., were not detected and thus, do not appear to impact net biogeochemical trends substantially (Fig. S2A; 2B).

### 4.3 Biogeochemical implications

Concerted research efforts have gone into understanding how Si availability impacts diatom metabolism. It is well established that low Si concentrations limit cell growth and reproduction which can then limit the extent of diatom blooms (Durak et al., 2016), but Si as a limiting factor or co-limiting factor for primary production is less understood (Maniscalco et al., 2023). Maniscalco et al. (2023) found that rather than Si stress alone inhibiting primary production, it can be co-limiting with N when both N and Si are depleted. In our case, NO_3_^-^ concentrations remained >4 µM and were not considered limiting to growth, potentially suggesting Si alone can independently limit diatom primary production. Concurrent measurements during the NA EXPORTS cruise did not find a strong response by diatoms to NO_3_ ^-^ addition, further suggesting nitrate was not independently limiting (B. Jenkins, pers. comm.). Given the key contribution of diatom production to total rates in this system (Meyer et al., 2024), the extremely low Si(OH)_4_ concentrations (average of 1.0 ± 0.5 µmol L^-1^) and abundance of light and other nutrients suggests Si availability was the primary control on diatom NPP during PI. Our gene expression data supports this physiological observation with variable silicon transporter expression corresponding to changes in ambient silicic acid concentrations. Although further research into Si limitation as a community-wide control on diatom growth in open-ocean environments is still warranted.

During the decline phase of the NA spring bloom, there was a transition from a diatom-dominated community to one dominated by other, non-diatom phytoplankton taxa. These other phytoplankton groups (particularly smaller celled groups such as haptophytes) can often be responsible for a majority of the primary production and nutrient cycling in various regions both within and outside of bloom conditions (Juranek et al., 2020; Meyer et al., 2022; Meyer et al., 2024). Our results support previous findings (Sieracki et al., 1993) which suggest that as diatoms decline upon Si(OH)_4_ depletion in PI, haptophytes capitalize on reduced resource competition and increase their gene expression related to new production. This creates a better balance between the contribution of small (i.e., non-diatom) and large (i.e., diatoms) cells to total community uptake trends in PII. However, the role of diatoms in NPP trends is still important, as suggested by the strongest relationship between diatom *NRT*:*Psb* and NPP by the large size-fraction across both PI and PII (R^2^ = 0.63; Fig. 6A). On the other hand, dinoflagellate and haptophyte trends between *NRT*:*Psb* (Fig. S4; See Supplemental Results S2.4.) and NPP by the large size-fraction are slightly lower at R^2^ = 0.62 and 0.53, respectively. While this trend is likely specific to Si(OH)_4_ limiting conditions, as it will disproportionally limit diatom growth and productivity, it represents larger ecosystem dynamics with diatoms appearing to respond more readily to changes in environmental conditions relative to haptophytes and dinoflagellates.

Interestingly, there was a strong and significant positive relationship between the relative proportion of diatoms and haptophytes in the surface community and sediment traps (Kramer et al., 2025), further supporting the notion that these two phytoplankton groups are disproportionally important to carbon export and biogeochemical cycling.

Our findings suggest frequently occurring storms throughout the spring in the North Atlantic can have a profound influence of spring bloom dynamics, community composition and phenology. Much uncertainty exists regarding how climate change will impact storm events in the North Atlantic (Wolf et al., 2020), making it difficult to assess whether the findings presented here are exceptions or will become more frequent in coming years. Understanding how different phytoplankton groups respond to these storm events, particularly in terms of variable mixed layers and nutrient concentrations during the bloom, and how the interplay of resource competition impacts ecosystem biogeochemistry is critical to understanding carbon export potential and future climate change in the region.

## 5. Conclusions

Our study provides new insights into some of the molecular underpinnings that control bloom dynamics in this climatologically important region and suggests that diatoms and haptophytes are dominant drivers of primary production in this system. High rates of primary production, primarily diatom driven, appear to have controlled Si(OH)_4_ depletion during the North Atlantic annual spring bloom, resulting in sustained Si(OH)_4_ limitation in the diatoms. As the bloom declined, Si(OH)_4_ depletion creates a decoupling between silicon metabolism from community carbon and nitrogen metabolism, as shown through the activity of two key silicon transporters relative to key photosynthesis (*Psb*, *Rbcl*) and nitrogen metabolism genes (*NRT*, *AMT*). Due to their lack of silicon requirement, haptophytes also appeared to metabolically respond to decreased diatom productivity and apparently reduced competition for resources during the second phase. Diatoms remain present in the system and are responsible for net ecosystem dynamics, particularly carbon export, despite the increased contribution of haptophytes during bloom decline. These findings support previous studies which suggest diatoms are responsible for net bloom dynamics while highlighting the key role haptophytes play during bloom succession over time.

## Data Accessibility Statement

All data presented here are available at the SeaWiFS Bio-optical Archive and Storage System (SEABASS; seabass.gsfc.nasa.gov/cruise/EXPORTSNA) or at the National Center for Biotechnology Information (NCBI; ascension number PRJNA1072555).

## Acknowledgements

We thank the EXPORTS leaders, D. Siegel and I. Cetinic, the chief scientists, D. Steinberg, J. Graff, D. Siegel, and C. Lee, and the captain and crew of the RRS *James Cook.* We also thank Yuanyu Lin, Emily Speciale, and Natalie Cohen for bioinformatics support, and Colleen Durkin for helpful comments on the manuscript.

