## Supplemental Information for "Phytoplankton phenology through gene expression during the North Atlantic spring bloom decline"

The supplementary information includes:

Supplementary Methods

Supplementary Results

Figures S1 – S7

Legends for Tables S1 – S3

**1. Supplemental Methods**

*1.1. KEGG Pathway Module Analysis*

The two CPM datasets were used to generate heatmaps of CPMs by day and hierarchical clustering dendrograms of KEGG pathway modules assessing relatedness (via Euclidean distance) of pathways through time or sample day characteristics based on metabolic pathway trends. After standardizing the CPM data, these plots evaluate relatedness on a -3 to 3 scale with 3 indicating strong, positive relatedness and -3 indicating strong, negative relatedness. The two plots highlight how differences in the normalization lead to varying statistical relatedness amongst pathways and how certain pathways may be more related within a specific phytoplankton group relative to what is preserved at the community level. Figures were generated using the clustergram function in MATLAB 2024b.

**2. Supplemental Results**

*2.1. Dinoflagellate and Chlorophyte Taxonomic Representation*

Major dinoflagellate genera include *Symbodinium*, *Alexandrium*, and *Heterocapsa*, at 33%, 27%, and 14% of total reads, respectively (Fig. S1). Major chlorophyte genera include *Micromonas* (48%), *Tetraselmis* (24%), and *Bathycoccus* (15%; Fig. S1).

*2.2. Dinoflagellates and Chlorophyte Transcript Abundances*

The most variable transcript abundances for any dinoflagellate-associated genes using either normalization method are in the photosynthetic gene *Psb* which increased substantially, from YD 126 to YD 142 (Fig. S2A, 2B). Moreover, the trends between phases differed by an order of magnitude with the PI average for *Psb* at 792 ± 389 whereas the PII average was 9,775 ± 3,335 transcripts. Other photosynthesis, nitrogen and vitamin metabolism genes also displayed variability between phases, however, no trend was as apparent as *Psb*. In fact, B_12_-dependent methionine synthase (*METH*), B_12_-independent methionine synthase (*METE*), *NRT*, *NR*, *NiR*, and *AMT* all decreased from PI to PII. Interestingly, *Rbcl* transcripts were not detected in dinoflagellates.

In both transcript abundance normalizations, chlorophytes exhibit some differential patterns compared to the other phytoplankton groups, but not as strong as those detected in diatoms and haptophytes between phases (Fig. S2A, 2B). While transcript abundances were higher in the haptophytes, the chlorophyte expression of *METH* is noteworthy as it is one of the genes with the highest transcripts. The expression of *IDH* is similar; *IDH* transcripts were higher for dinoflagellates, but they are proportionally much higher relative to the other gene abundances for chlorophytes. Chlorophytes have high transcript abundances for arginase (*Arg*), which is involved in the urea cycle and ranged from 35.8 – 256 CPM, with peak counts coinciding with YD 128. Interestingly, chlorophytes are the only phytoplankton group to have higher average transcript abundances for Photosystem I (*Psa* with 31.2 CPM) relative to Photosystem II (*Psb* <1 CPM). Additionally, many of the genes highly expressed in diatoms, dinoflagellates, and haptophytes, including *NRT*, *AMT*, and *Rbcl*, were not detected in chlorophytes.

*2.3. KEGG module trends*

Module plots can also be used to evaluate relatedness amongst sample days based on KEGG module trends (Fig. S3). Here, we see interesting trends in which sample days are considered the most related or disrelated based on physiological characteristics. The strength of these relationship (0 being low and 3 being high) reflects correlations between the temporal expression patterns of genes involved in each of these processes. Unsurprisingly, YD 128 appears the most distinct within the diatom group. This is consistent with the highest transcript abundances for photosynthesis, vitamin, nitrogen, and silica metabolism occurring on this day as well as the highest overall diatom transcripts. YD 126 appears the most unique for haptophytes (Fig. S3). This is likely for the opposite reason as for the diatoms as this is a period of less overall transcription by haptophytes. YD 132 branches independently for chlorophytes, a trend that appears driven by transcript abundances of isocitrate lyase, malate dehydrogenase, and cytochrome b6-f complex. Within the dinoflagellate modules, trends by sample day appear overall more homogenous with no single YD branching independently of the others.

*2.4. NRT:Psb ratio trends*

*NRT*:*Psb* ratios were calculated for diatoms, dinoflagellates, and haptophytes (Fig. S4), assuming *NRT*:*Psb* ratios would broadly correlate with the N and C uptake trends. These ratios showed substantial differences among groups with diatoms and dinoflagellates exhibiting similar ratios (averages are 0.24 ± 0.18 and 0.13 ± 0.22, respectively) but haptophytes exhibiting much larger ratios (average = 4.05 ± 2.27). Interestingly, the haptophyte ratio only drops below 1:1 on one sample day (YD 128) whereas the diatom and dinoflagellate ratios never reached 1:1 (Fig. S4). The haptophytes exhibit a stronger average ratio in PII (4.77 relative to 1.18 in PI) whereas the diatom and dinoflagellate ratios decrease from 0.34 to 0.21 and 0.48 to 0.04 from PI to PII, respectively (Fig. S4). Diatom and dinoflagellate *NRT*:*Psb* ratios still exhibited strong relationships to NPP in the large size-fraction with R^2^ = 0.63 and 0.62, respectively.

Additionally, a principle component analysis was run using MLD, macronutrient concentrations, total phytoplankton production (NPP, new production), large size fraction biomass (Chl *a*, PC, PN), and gene and uptake ratios of Si(OH)_4_ (Fig. S5). While this analysis explained most of the variability (93.6%), MLD, macronutrient concentration, NPP, and new production all clustered independently from the large size-fraction biomass and Si(OH)_4_ ratios which clustered very close together along components 1 and 2 (Fig. S5). The closest of all relationships is between Si4:*Psb*, Si4:*NRT*, and large size fraction *ρ*Si:*ρ*DIC, suggesting that both Si4 and Si7 gene expression may be promising bioindicators of Si(OH)_4_ limitation.

The ratio of haptophyte *NRT*:*Psb* remains consistently high through the final day of our observation period (Fig. S4). This is also similar with the trends in size-fractionated NPP and new production through YDs 145-149 which only varied by 2.31 - 4.03 mmol C m^-2^ d^-1^ (Meyer et al., 2024). Overall, whether this period represents the typical bloom succession and a return to lowered NPP dominated by the haptophyte community is uncertain. However, we note that Chl *a* concentrations from the PAP mooring indicate that surface concentrations recovered to values >1.5 µg L^-1^ by mid-June 2021 (~one month after our observation period ended) (data available at projects.noc.ac.uk/pap/data/pap-april-2021).

**Supplemental Figures**

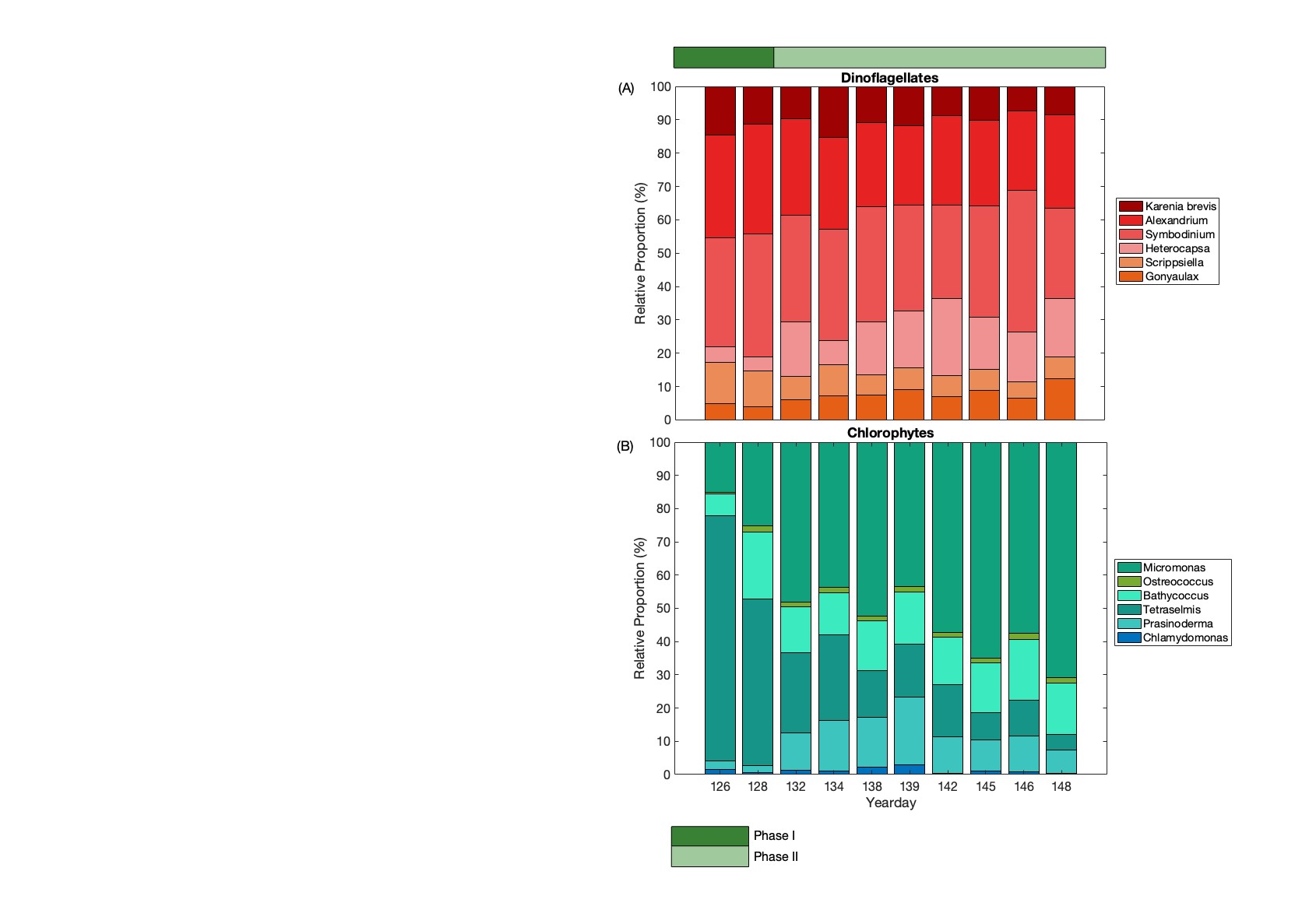

**Supplemental Figure S1.** The relative percentage (%) of the most abundant genera in dinoflagellates (A) and cryptophytes (B) based on sequence reads.

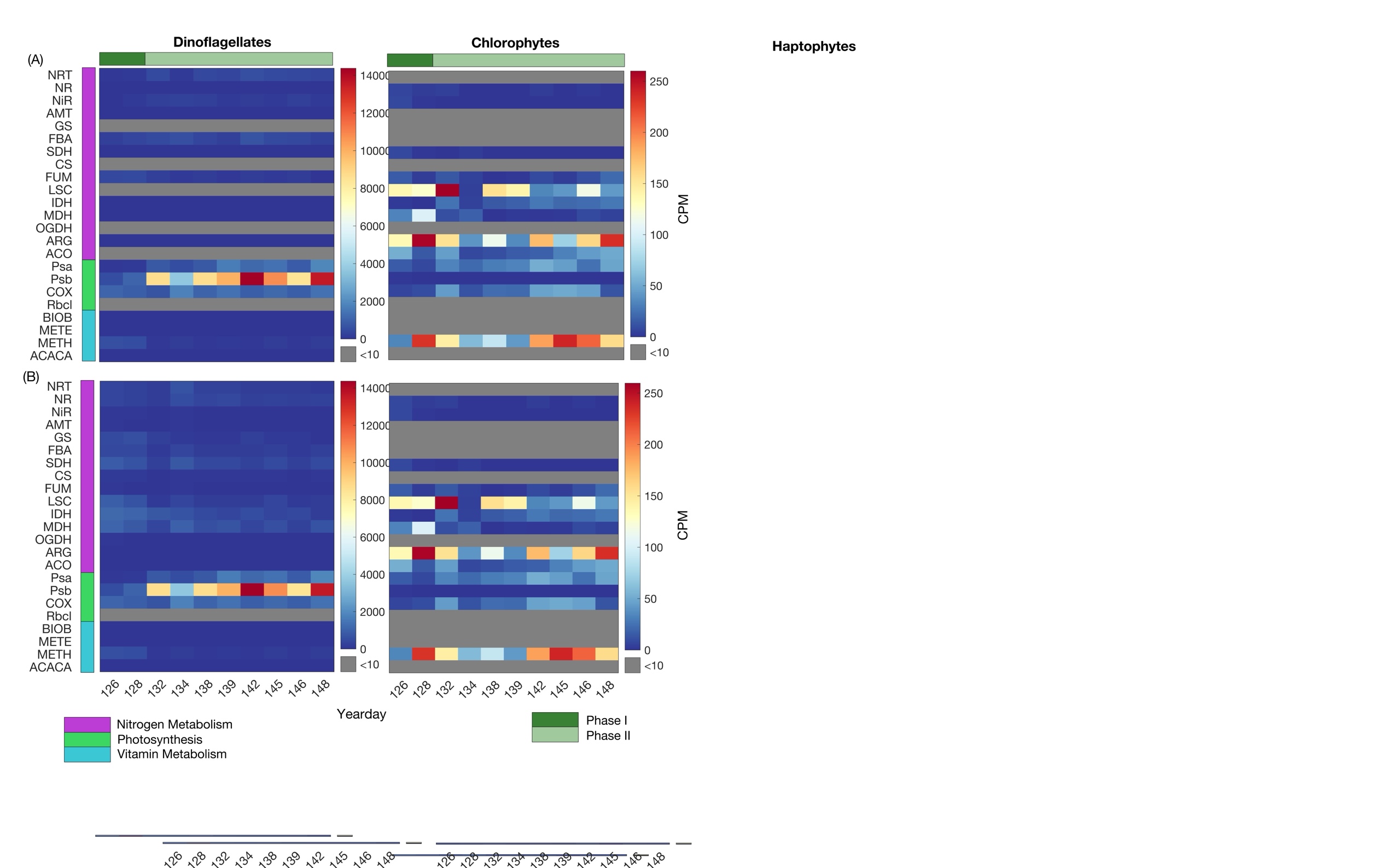

**Supplemental Figure S2.** Heatmaps of transcript counts per million (CPMs) of key nitrogen metabolism, cell maintenance, photosynthesis, and vitamin metabolism genes (indicated by left hand bars) for dinoflagellates and chlorophytes with counts normalized within each taxonomic group (A) and to the whole community (B). Plots are by yearday with the color bar indicating CPM for each gene. Green bars indicate phase. Full gene names are: nitrate transporter (NRT), nitrate reductase (NR), nitrite transporter (NiR), ammonium transporter (AMT), glutamine synthase (GS), flavodoxin (FBA), succinate dehydrogenase (SDH), citrate synthase (CS), fumarate (FUM), succinyl-CoA synthetase (LSC), isocitrate dehydrogenase (IDH), malate dehydrogenase (MDH), oxoglutarate dehydrogenase (OGDH), arginase (ARG), aconitase (ACO), photosystem I (Psa), photosystem II (Psb), cytochrome c oxidase (COX), rubisco (Rbcl), biotin synthase (BIOB), methionine synthase (METE), methyltransferase (METH), and acetyl-CoA carboxylase (ACACA).

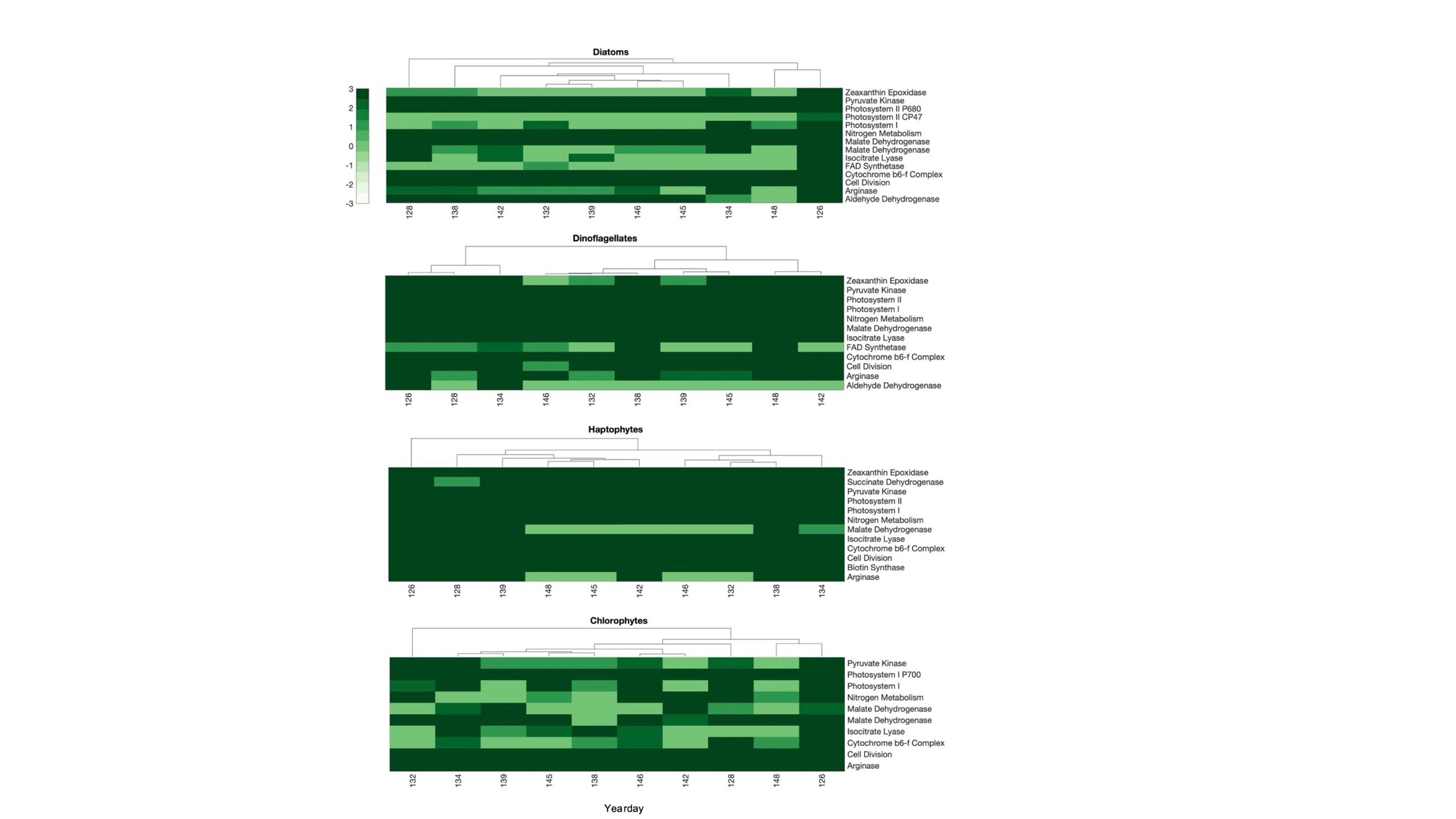

**Supplemental Figure S3.** Clustergrams of Julian day relatedness based on the relationships between KEGG pathway modules, by phytoplankton group. CPMs driving the module relationships were normalized within each phytoplankton group. Branch length on the dendrogram indicate relatedness based on Euclidean distance. The color bar indicates the strength and directionality of the relatedness of modules.

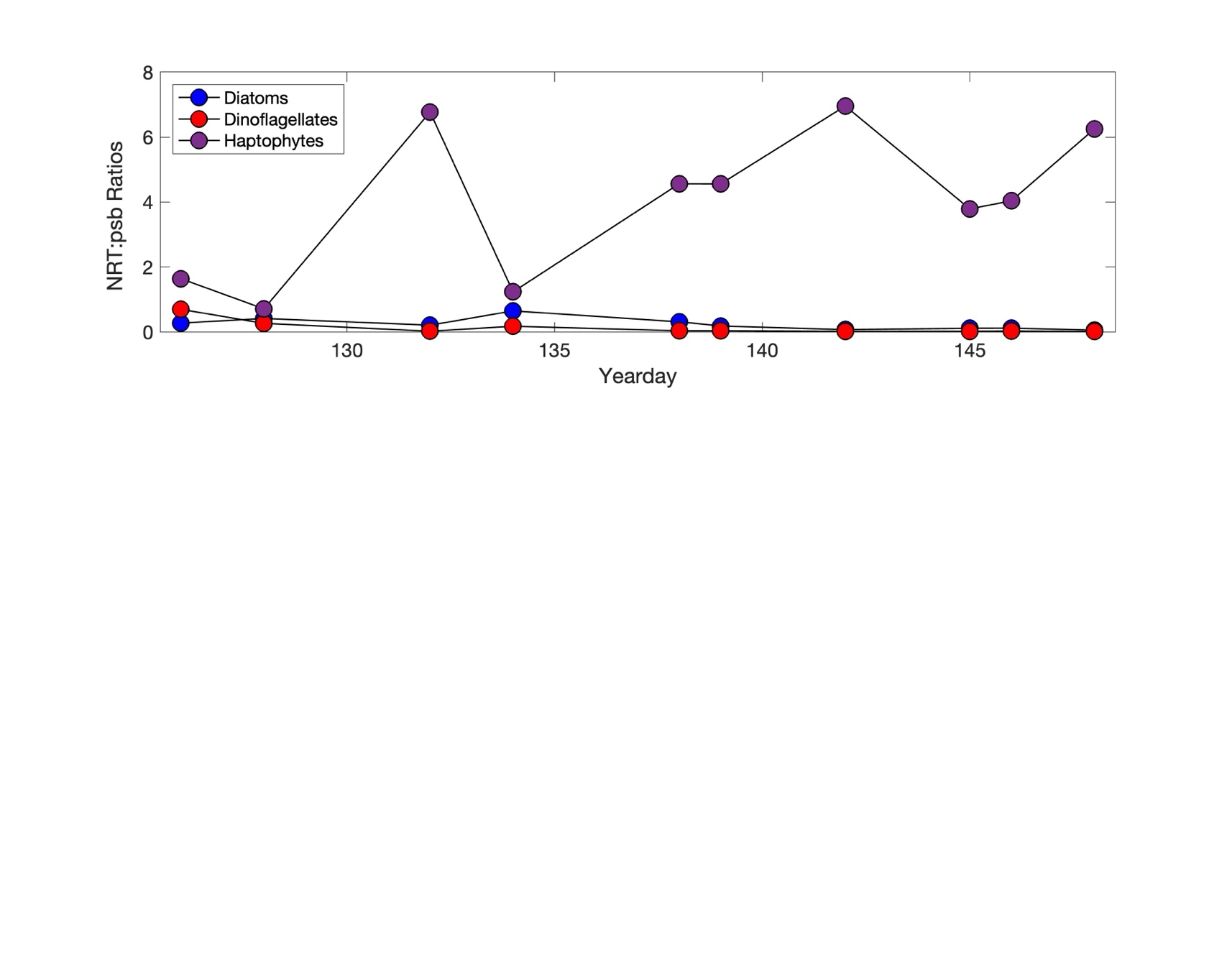

**Supplemental Figure S4.** Ratios of nitrate transporters (*NRT*) to Photosystem II (*Psb*) for diatoms, dinoflagellates, and haptophytes by yearday.

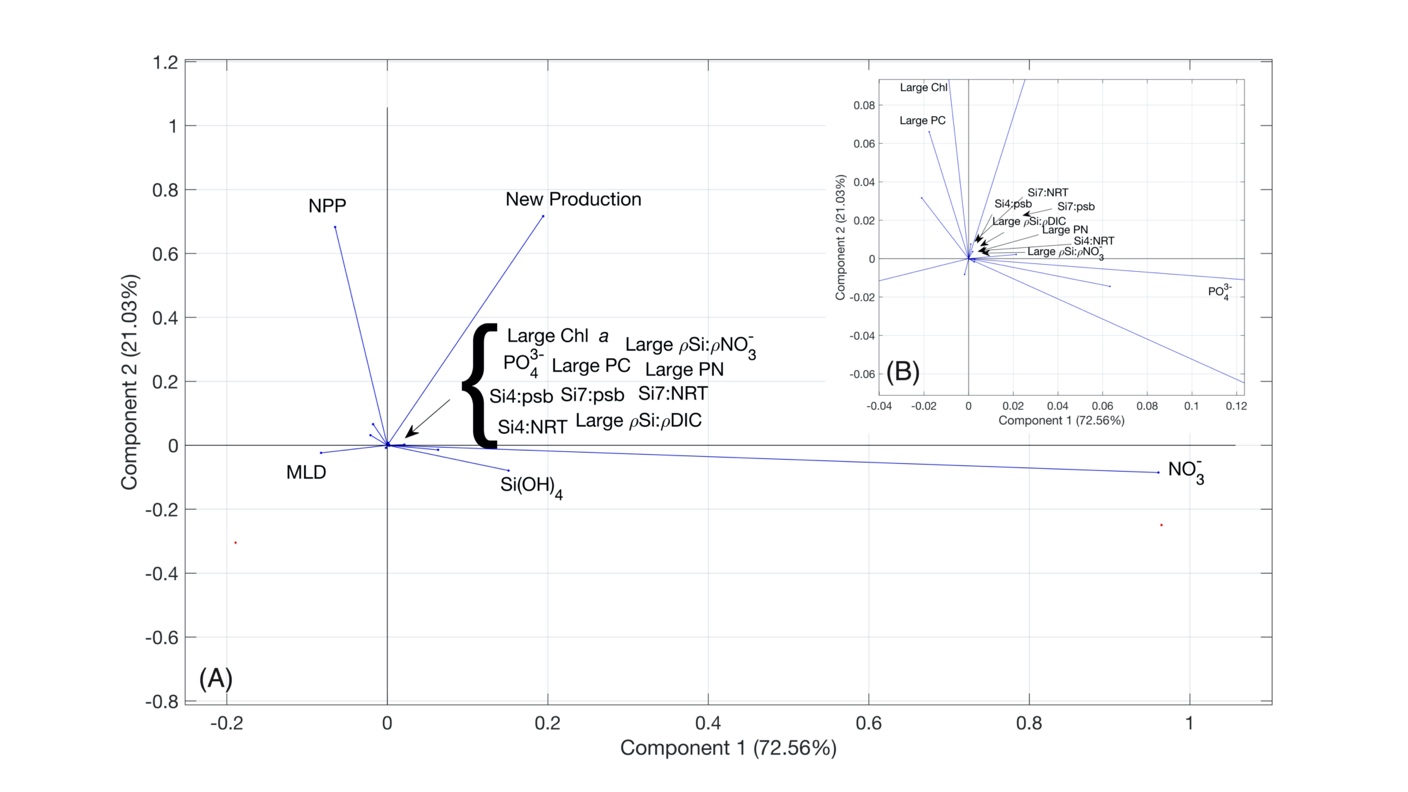

**Supplemental Figure S5.** Principle component analysis relating environmental parameters, production parameters, and size-fractionated phytoplankton biomass parameters with physiologically interesting gene ratios and Si, C, and N isotopic uptake ratios. The inset map (B) is a zoomed-in version of the full PCA to show greater detail amongst parameters that cluster very closely together.

**Supplemental Tables**

**Supplemental Table 1**. Source information for the 7 Si-related genes that appear in our metatranscriptomic sequences. All information shown here comes from the National Center for Biotechnology Information (NBCI) and can be found via each FASTA file’s accession number.

| **ID** | **Source Organism** | **% ID** | **E-value** | **Source** | **NCBI Accession ID** |
| --- | --- | --- | --- | --- | --- |
| Si1 | *Chaetoceros muelleri* | 95.67 | 0.0 | Encinas-Yanez et al., unpublished | UAD82345.1 |
| TFSGINTSRIQATSTNKNMSSCRVLKQIYSTVLLIFSVVTIMGLIAQRQTRISADFHPILAYIVIWISIVWLSMIEGSQGSLVGLSSINADLYRDSHPIAYKCTAITNQGNNINRYLIGRQFMVVLVVFCVNLCGGPLESASGIWNYPNWVNAIFLTVGFAMILFTCMVGQLNSQVNASLSMLDYINNYFALVTVWVALGIEYSGLLHSSYLLQMFVTYLSDRTIDERTKASSRSLSWAGKVFFWTRCIMSLAILIFCFAVTLEALFSGKTTMWEGVPPGSAVVVFFILMSIVGMLEGMQIAFFAVAKIPAAERGNSTFALKTCNLLFQGGEDGTNNLPQFMIGRQLFVVTCMFFIARVTSIQIPQGEYAQGDQGRPSANNNNNIFGVSDTVQEILMNSGLLGALITTTVGSLAWQLIASAFPLSFLANPLTYALLRVCLFLEATGICSGAWVLANIHQRVAGFQRDEVYIGTAEDRAKREMGDDTDHFPIGPGYIVKLPGFAHHAPKSLKELLATTNGGSYQFDELYIFPENDSSSSSFSEPGNSSFDGHDEEENISSTSSLVKPLHNTNLIPTEPLDEGSSLNNTRMHDSEYSQEEIEEEEEAGKESDLSTTRRIIHQDTTLEQNEAIQSSKVNNF | | | | | |
| Si2 | *Phaeodactylum tricornutum* | 66.25 | 0.0 | Bowler et al., 2008 | XP_002184375.1 |
| SLYLETAFRWKISYRGLKSNNLATSLLVIVFTAVKENKMTDGSISAFTLLKNLYSVCLLIFSVVIVMGLIFSGQTKIASEVNPWVAFIVIWAALLWLSMVEGGQGALVGLAPINFDLYKDTHPKTYMSTKIAHVGDNLDRYLLGRQFMVLFIVFSTNMAGAPLPNSQLWGFPDWVITIFLGTGIAMILMTNMIGQLNTQVNAAHCMLDYINSYFAVFTFYVAMAIEFSGLLHSSYVIQIIVGLMAGKNIESNEPPRSGFVLLFFWLRCLASIAILFFCLAVTIEALFAGQTTMWEGVPNGLAVVLFFVLMSVVGLLEGMQIAFFAVTKLRESERGSGVFAKRTCELLFRGDGHNLPGFMIGRQITVTLNFFVIARVTSLNIVPGKGENVFGVSDAMQEFFNTGLLGALITTILASIAWQLVASAFPIAFLSNPIVYVFLVLALLLEGTGVCNGAWVLAAIHKQIAGYQRDEVYIGTAEERAQRNMEDKPVAAGAGHIFKLPGFVDMPEALEALMLNDPTVAEFIRKISQHSELEMVEEESRL | | | | | |
| Si3 | *Cylindrotheca fusiformis* | 100.00 | 0.0 | Hildebrand et al., 1998 | AAD13804.1 |
| MVSVIDGIKQFYSMALVIFSVVIVTALMFTDNTKLAKDAHPVAALVIMWLGILWMSMVEGGQCSMVGLPPIDRDLYKESHPITYKICSLGHKGNNLDRYLMGRQFMVIFINFTINLCGAPLEGAEVLGLPEILTDIFLGSGIAMVLTVVTIGQLTAQVNASHCMLDYINTHFMTFTLYVTLVIEATGVMHSCYLIRDMFYHAAGKPVETNEPPRSAVQNLFHWGRVVFSLGVLCFALAVTIEALFNGKTTMWEFIPNGVAIVLFILLMSVVGLLEGMQIAFFAVAKIPKAERGDHPFARKTCELLFKGKGRNLPGFMVGRQMTVTLCFFIIARVTTLDIEVGVDDNIFGVSDGIQEFFNLGFLGAIITTILASIAWQLVASAFPIAFLSNPIVYIVLRIVLLIEATGICAGAWFLGMIHKKVAGFQLDEVYVGTAEERAAGMKPDHSIHAGREFTMGTNVLNDRKNWEETIANLSAKETFSVRRERMLKNIRELRAMAEEASSPEEKATFETALTMETKALNKLNEEQEKEATLQKDSSDTENEADMA | | | | | |
| Si4 | *Fragilariopsis cylindrus* | 81.45 | 0.0 | Mock et al., unpublished | OEU10496.1 |
| THTTTKHTMADHSHGTNTTWDWIKIAYSCFLVGFSVVVVLAXIFTGNTQLARDIHPIAAAIILWVLIIWMSMIEGGQCSMVGLPPIDRELYRESHPITYKITGWGHKGDNLDRYLMGRQFMVIFVNFTIGLCGAPLDSTVSVMGLPPWIIAIFLGSGIAMVLQNVTIGQLTSQVNASHCMLDYINTHFMTFTYFVAFTIEATGVMHINYVIRMMAYYAAGKPVESNEPPKEGLTLLFFWGRVLWSAGILVFAIAVTCEALFRGKTSLWDGVPEIIGLVLFFVLMSAIGLLEGMQIAFFTVSNIPKSERGNSSLALKTCHILFKNGGKNLPGFMCGRQITVTLCFFIIARVTTINVAIGEEPNIFGVADWVQKMFNLGFMGAITTTILGSVAWQLVASSFPIAFLSNPIVFVFLQAALFLEATGICAAAWFLALLQKKVMGYQYDEVYVGTPEERAAKDHADDNEAQMHLDMGTNVLAPGINDAVGFNSNWAEGDYTDRRNQTLQNIAELRLQINLCVTEEEKEAFQMALKLEVQQLERTNKEQETSVRNLGKGGDAEAV | | | | | |
| Si5 | *Fragilariopsis cylindrus* | 80.42 | 0.0 | Mock et al., unpublished | OEU11729.1 |
| MLTEDTSRNSSSKGIFMWFKEIYSTCLLIFCSIIVLTVIFEKNTKLSESSSWAAFFVFWVALYWLSMVEGGQASLVGLPPVDMELYKDSHPTTHKIMKVINRGDTLDRYLMGRQFMVLALVFVENLCAHANDSSLPILGMPIVINKIFLDTGLAVFFMTAMIGKISAQVNASRCMLDYVNNFFAYFTMQTARLIEVSGLLHCCYPVQIFFARAAGQPLESKEPPRSLVQNIFFWSRVLLSTAILAFAFAVTLYAIVNGQTTMWEGVPSWLTIVLFFLFMSVVGMLEGMQIAFFAIARMTEEERAKSTWAKKTCDLLFEGDGRNLPGFMIGRQMCVTLCFFIIARVTTVQLQPGDENVFGVSDATQAFFATGLLGALITTIVASIAWQLVASAFPMAFLSTPITYILLRFCLGLEWTGICQGSWVVARLHRKIVKFKRDEVYIGTAEERAAKAKANPTKFEGDHEDAMNVKPGHMYPGVPTLPPDFKGVNRTLQEIEELESDLKEKQQEIEDRLTDLAAQKKRMLGVGEEC | | | | | |
| Si6 | *Skeletonema costatum* | 100.00 | 0.0 | Thamatrakoln et al., 2006 | ABB81824.1 |
| MTDAEIKKEELADAHDVELTPFTIFRYTYSVILLIFSIVLVVSLMFTGNTKLAADASPWAALFVCIAAVVWLSMIEGQQASLVGLPPVDPELYKETHPVTYLNAATAFLGDNLDRYLMGRQFMVLLVVFIINLCGAPSSGDADVLGMPGWLKTIFLDVGLGMIIFTCQLGQLTTQVNASHCMLDFINNYFALFTLYTAMCIEFSGVMHSSYLIQNVLSFASGKPIHSNEEPKRGFTLLFFWGRVLMSLAILGFSLAVVISALFQGRTMMAVKYPSVSNGASVFLFFFLMCIVGMLEGMQIAFFAVAKLPASERGTTFFGRKTCDLLFKGNGQNLPGFMIGRQLTVVASFFIVASITSMNIQPGNEDGNIFGVSDGAQAFLNLGFHAAVITTILASITWQLAASAFPIAFLNNPVTYVLLVFALFLEWTGLCAGAWVLARVMKKALKYEYDEVYVGTPEERAANNHADKDFADDTGKMYGGGFRGHAVGSHDALDGPIASKDEVEEEAV | | | | | |
| Si7 | *Phaeodactylum tricornutum* | 53.47 | 0.0 | Bowler et al., 2008 | XP_002183269 |
| YSTIALIFSVVCIHGLIFTQQTGLSENNSIIAVVVLWAAIFWLAMIEGGQASHVGLAPIDENLYKDSHPTTYKISAACNKGDNLDRYLLGRQFGVIFVVFCVNLSGGPIGGASLWGLPXIVQQIFFSTGFAMILFTCMVGQLNTQVNASICMIDFINNYVSLFTFWCCMVVEFVGIVHSSYAIQMAIGSMAGXPIZSKEPPKEGFTLMFFWLRVLWSFGLLGFSLXVTIAALFEGKTTMWDGVPEIXSLILFFGLMSVVGMLEGMQIAFFYVAKLSEEERGSNSWANRTCDLLYNKRNGLNLPGFMIGRQLSVVTCFFVVARVTTQSXEDGETNVLGLPDPVQNFLNFGFQGALITTILGSITWQLVAAVFPVAFLSTPITYILLRICLFFEWTGICNGAWVIAGILKSLFGYQLDEVYIGTPEERKARGMKDDSQREIAIPALSGFNGLPNFHSAPKSLRDLAESDPEVGAYLSDVAGTMKTGAPAAAAAPAAASYAEEDDEVEC | | | | | |

**Supplemental Table 2**. RNA sequence samples with their number of reads, percent GC content, and N50.

| **Sample ID** | **Number of Reads** | **%GC** | **N50** |
| --- | --- | --- | --- |
| 1-2-1A | 21864628 | 64.3 | -- |
| 1-2-1B | 18525876 | 63.6 | -- |
| 1-2-1C | 25267987 | 62.0 | -- |
| 1-4-1A | 28897299 | 64.5 | -- |
| 1-4-1B | 31340014 | 65.7 | -- |
| 1-4-1C | 22069239 | 62.2 | -- |
| 2-2-1A | 27132394 | 67.5 | -- |
| 2-2-1B | 28421928 | 68.1 | -- |
| 2-2-1C | 27601814 | 66.9 | -- |
| 2-4-1A | 32138506 | 67.2 | -- |
| 2-4-1B | 33106563 | 66.1 | 378 |
| 2-4-1C | 24957299 | 63.8 | 336 |
| 2-8-1A | 29151655 | 66.9 | 373 |
| 2-8-1B | 28283047 | 65.2 | 371 |
| 2-8-1C | 30095190 | 67.3 | -- |
| 2-9-1A | 27832681 | 66.7 | 360 |
| 2-9-1B | 25640012 | 66.3 | 356 |
| 2-9-1C | 32125981 | 65.1 | 370 |
| 3-2-1A | 31865420 | 65.6 | 378 |
| 3-2-1B | 28197340 | 67.2 | 380 |
| 3-2-1C | 27303715 | 67.9 | 385 |
| 3-5-1A | 29390093 | 66.9 | -- |
| 3-5-1B | 27814830 | 65.1 | 373 |
| 3-5-1C | 26706048 | 67.0 | 371 |
| 3-6-1A | 25387077 | 67.2 | -- |
| 3-6-1B | 31262047 | 64.9 | 429 |
| 3-6-1C | 29451453 | 64.7 | 400 |
| 3-8-1A | 28608270 | 66.3 | 394 |
| 3-8-1B | 29187154 | 66.5 | 375 |
| 3-8-1C | 25694969 | 68.4 | 367 |

| **YD** | **MLD (m)** | **ML Integrated PAR (mmol m^-2^ s^-1^)** | **NO_3_^-^ (mmol m^-2^)** | **PO_4_^3-^**  **(mmol m^-2^)** | **Si(OH)_4_**  **(mmol m^-2^)** | **GPP**  **(mmol C m^-2^ d^-1^)** | **NPP**  **(mmol C m^-2^ d^-1^)** | **New Production**  **(mmol C m^-2^ d^-1^)** | **Small Chl *a***  **(%)** | **Large Chl *a***  **(%)** | **Small PC**  **(%)** | **Large PC**  **(%)** | **BCD (mmol C m^-2^ d^-1^)** |
| --- | --- | --- | --- | --- | --- | --- | --- | --- | --- | --- | --- | --- | --- |
| 126 | 42 | 6.48E+05 | 140.4 | 8.4 | 5.6 | 561.2 | 146.6 | 99.36 | 19.1 | 80.9 | 41.8 | 58.2 | 10.2 |
| 128 | 33 | 6.39E+05 | 144.6 | 8.0 | 3.0 | 274.3 | 93.9 | 65.74 | 16.2 | 83.8 | 43.2 | 56.8 | -- |
| 132 | 43 | 6.04E+05 | 330.2 | 21.1 | 73.6 | 558.4 | 89.9 | 133.5972 | 54.0 | 46.0 | 61.9 | 38.1 | -- |
| 134 | 19 | 6.70E+05 | 364.0 | 24.4 | 40.7 | 231.5 | 45.9 | 49.5495 | 28.9 | 71.1 | 48.6 | 51.4 | 6.8 |
| 138 | 64 | 7.66E+05 | 192.8 | 12.4 | 22.6 | 355.2 | 95.4 | 65.49906 | 46.9 | 53.1 | 64.0 | 36.0 | 22.3 |
| 139 | 28 | 1.75E+05 | 156.8 | 9.8 | 9.6 | 201.3 | 74.5 | 49.25448 | 41.1 | 58.9 | 52.2 | 47.8 | 14.9 |
| 142 | 74 | 7.58E+05 | 199.4 | 13.3 | 45.8 | 279.6 | 48.4 | 20.19534 | 40.6 | 59.5 | 59.0 | 41.0 | 23.3 |
| 145 | 59 | 6.52E+05 | 132.2 | 9.0 | 24.2 | 150.7 | 56.6 | 30.60486 | 47.4 | 52.6 | 59.2 | 40.8 | -- |
| 146 | 23 | 2.35E+05 | 177.1 | 11.9 | 54.1 | 177.5 | 34.4 | 16.35612 | 37.7 | 62.3 | 53.4 | 46.6 | 14.9 |
| 148 | 31 | 8.40E+05 | 134.8 | 8.7 | 25.0 | 196.5 | 58.6 | 32.23176 | 47.7 | 52.3 | 36.7 | 63.3 | 10.1 |

**Supplemental Table 3**. EXPORTS North Atlantic environmental parameters, production parameters, phytoplankton biomass parameters, and bacterial carbon demand (BCD) by yearday (YD) within the mixed layer. Environmental parameters include mixed layer depth (MLD), mixed layer integrated photosynthetic active radiation (PAR), nitrate (NO_3_^-^), phosphate (PO_4_^3-^), and silicic acid concentrations (Si(OH)_4_). Production parameters include gross primary production (GPP), net primary production (NPP), and new production.

**Supplemental Table 4.** Table of Pearson correlation coefficients (R), R-Squared (R^2^), and p-values between large size-fraction uptake ratios (*ρ*Si:*ρ*DIC and *ρ*Si:*ρ*NO_3_^-^) vs. gene ratios (Si4:*Psb*, Si7:*Psb*, Si4:*NRT*, Si7:*NRT*) through the whole observation period.

|  | R | R^2^ | p-value |
| --- | --- | --- | --- |
| Si4:*Psb* vs. *ρ*Si:*ρ*DIC | -0.60 | 0.36 | 0.16 |
| Si7:*Psb* vs. *ρ*Si:*ρ*DIC | -0.81 | 0.65 | 0.10 |
| Si4:*NRT* vs. *ρ*Si:*ρ*DIC | -0.75 | 0.56 | 0.30 |
| Si7:*NRT* vs. *ρ*Si:*ρ*DIC | -0.86 | 0.74 | 0.22 |
| Si4:*Psb* vs. *ρ*Si:*ρ*NO_3_^-^ | -0.45 | 0.20 | 0.25 |
| Si7:*Psb* vs. *ρ*Si:*ρ*NO_3_^-^ | -0.71 | 0.51 | 0.43 |
| Si4:*NRT* vs. *ρ*Si:*ρ*NO_3_^-^ | -0.57 | 0.32 | 0.29 |
| Si7:*NRT* vs. *ρ*Si:*ρ*NO_3_^-^ | -0.71 | 0.51 | 0.59 |
